# A Role for Visual Areas in Physics Simulations

**DOI:** 10.1101/2021.09.14.460312

**Authors:** Aarit Ahuja, Theresa M. Desrochers, David L. Sheinberg

## Abstract

To engage with the world, we must regularly make predictions about the outcomes of physical scenes. How do we make these predictions? Recent evidence points to simulation - the idea that we can introspectively manipulate rich, mental models of the world - as one possible explanation for how such predictions are accomplished. While theories based on simulation are supported by computational models, neuroscientific evidence for simulation is lacking and many important questions remain. For instance, do simulations simply entail a series of abstract computations? Or are they supported by sensory representations of the objects that comprise the scene being simulated? We posit the latter and suggest that the process of simulating a sequence of physical interactions is likely to evoke an imagery-like envisioning of those interactions. Using functional magnetic resonance imaging, we demonstrate that when participants predict how a ball will fall through an obstacle-filled display, motion-sensitive brain regions are activated. We further demonstrate that this activity, which occurs even though no motion is being sensed, resembles activity patterns that arise while participants perceive the ball’s motion. This finding suggests that the process of simulating the ball’s movement is accompanied by a sensory representation of this movement. These data thus demonstrate that mental simulations recreate sensory depictions of how a physical scene is likely to unfold.

## Introduction

An intuitive understanding of the laws of physics is one of the fundamental, underlying aspects of our daily interactions with the world around us. Almost every action we take relies on our brain’s ability to effortlessly compute possible physical outcomes. Given the diversity of scenarios that we navigate every day, it seems likely that we use a variety of different strategies to parse physical scenes. One strategy that has recently risen in prominence is that of simulation (Bates et al., 2019; Fischer et al., 2016; Hegarty, 2004; Rajalingham et al., 2021; Ullman et al., 2017). Simulation refers to one’s ability to run a “mental model” of a physical scene in order to determine an outcome. Physics simulations have long been a centerpiece of industries such as computer graphics, video games, and even architecture and construction. The idea that the human brain might rely on simulation as a general mode of cognition is not new. Dating as far back as the 18^th^ century, philosophers David Hume and Adam Smith postulated that the way human beings enact empathy is by adopting mental models of one another (Hume, 1739; Smith, 1759). Kenneth Craik extended this concept beyond human interactions and argued that many everyday problems are resolved through the use of “small-scale models” that reflect reality while existing entirely in the mind (Craik, 1943). Most recently, the simulation theory has been put to the test in the lab in the context of physical scene understanding. For instance, Battaglia et al. have shown that computational models that make use of an “intuitive physics engine” to simulate rigid body dynamics of object interactions successfully capture human behavior on a variety of psychophysical tasks (Battaglia et al., 2013).

While the fact that people appear capable of physics simulation is an exciting finding, it marks only the beginning of potential interesting investigations into this topic. If we are to leverage our knowledge of simulations in the brain for real-world applications (for example, computer vision) or to inform therapeutic health interventions, it behooves us to fully understand the underlying mechanisms and circuits that enable our mental models. One immediate question that comes to mind concerns the fabric of the simulation itself. Are the physical simulations we employ represented in the brain as merely a series of computations between abstractly held entities? Or do our mental models evoke sensory representations of the physical interactions being simulated, even if they are not literally perceived?

Past research on the interplay between cognition and vision suggests that the latter might be true. For instance, decades of neuroimaging experiments have demonstrated that when we imagine stimuli with our eyes shut (a phenomenon termed “mental imagery” in the literature), early visual areas are activated as if those same stimuli were in fact being perceived (Klein et al., 2004; Kosslyn et al., 1995, 1997, 2001). A similar finding emerges when people are asked to rotate 3D shapes in their mind. Specifically, area MT, which is known for its role in the perception of motion, appears to be activated during mental rotation of objects, even in the absence of a visual stimulus (Shelton & Pippitt, 2006). Based on these discoveries, scientists have concluded that the process of internally envisioning stimuli that are not in fact present can evoke “simulated” sensory representations (Pearson, 2019). Given these findings, one may hypothesize that physics simulations of events could also evoke corresponding, internally generated visual representations.

We were interested in testing exactly this hypothesis. What might a visual representation of a physics simulation in the brain look like, and where might we expect to find its neural correlates? Considering the dynamic nature of our mental models, simulations of physical interactions must often incorporate the movements of objects, even if these movements are not being directly perceived at the time of the simulation. We thus theorized that, as in studies of mental rotation, we might find evidence of visual representations of physics simulations in brain regions such as area MT that are specialized for the perception of motion. This notion is further supported by past research showing that area MT can become active in response to static stimuli that contain elements of implicit or implied motion (Kourtzi & Kanwisher, 2000; Lorteijie et al., 2011).

In a previous study, we designed a novel task in which participants had to predict a ball’s likely trajectory as it fell through an obstacle filled display (Ahuja & Sheinberg, 2019). We refer to this as the *ball fall* task. The task could be solved by internally simulating the ball’s trajectory (including the various physical interactions encompassed within it) and using the outcome of that simulation to inform one’s answer. Through our prior work, we have provided behavioral, oculomotor, and computational evidence consistent with the idea that participants engage in simulation as they perform the ball fall task. In the present study, we recruited a new cohort of participants and asked them to perform the ball fall task while undergoing functional magnetic resonance imaging (fMRI). We sought to determine whether there was neural evidence supporting the idea that a physical simulation of an object’s trajectory would recruit neural circuits involved in the visual perception of that same trajectory. We hypothesized that if this indeed were the case, then we would observe voxel-wise activity pattern similarity in motion-sensitive regions of the brain between conditions in which participants simulated the ball’s motion trajectory, and conditions in which they perceived the ball’s motion trajectory. We found that this was indeed the case for all participants. This similarity effect between simulation and perception of the ball was not present outside of motion-sensitive brain regions. These findings provide evidence for a visual correlate of physical simulation.

## Methods

### Participants

Twelve individuals (4 male; 8 female) participated in this study. Participants were recruited from the Brown University campus and the surrounding community. All participants had normal vision and reported that they were not colorblind. Participants were screened for MRI contraindications, and were only included if they passed all screening requirements. Participants were compensated a base amount for their time, with additional compensation provided for correct responses on trials. Signed consent was received from all participants. The study was approved by the Brown University IRB.

### Motion Localizer

The first task the participants performed was a motion localizer task (Figure 3A). We used the motion localizer to define a motion-sensitive functional region of interest (ROI) for subsequent analyses. The localizer in this study was based on the one used in Sunaert et al., 1999. Localizer runs started with a 16-second lead-in period with only a yellow fixation point on screen with a black background. Participants fixated on the point for the entire 16 seconds. This was followed by randomly ordered 20-second blocks of white dots that either coherently moved in a given direction (i.e., the Motion condition), or remained completely stationary (i.e., the Static condition). During the Motion and Static conditions, the yellow fixation point remained on screen, and participants were required to continue fixating (while ignoring the white dots in the background). The white dots were presented in a circular area with a radius of 6 degrees visual angle around the yellow fixation point. White dots were 0.07 degrees visual angle in size and had a density of 69/degrees^2^. During the Motion condition, the white dots moved at 5 degrees/second, randomly changing direction once per second. Each run had 3 blocks of each condition (leading to a total of one minute per condition). Participants performed either one or two runs of the localizer task, depending on the time constraints of the session.

### Task

To evoke dynamic physics simulations, we used what we refer to as the ball fall task (Figure 1A). The stimulus used for a single trial in the ball fall task consists of one “ball” at the top, a set of semi-randomly arranged “planks” throughout the middle, and two “catchers” at the bottom. We refer to each display as a “board”. Participants were asked to determine which of the two catchers the ball would land in were it to be dropped from its given position, and indicate their choice by pressing one of two buttons.

**Figure 1:**
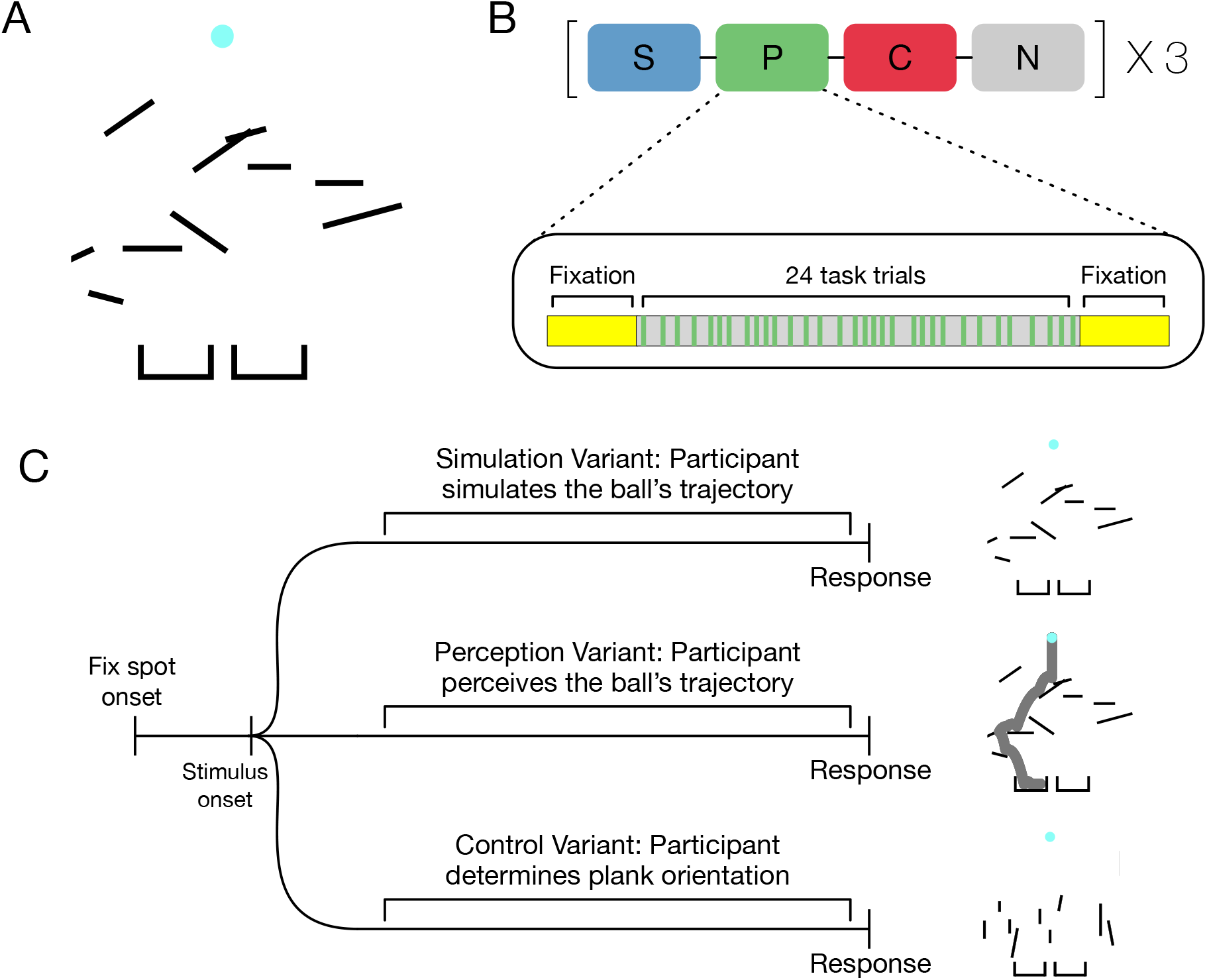
Task Design. (A) An example of a board that constituted the primary stimulus in the ball fall task. Participants had to determine which of the two catchers the ball would land in if dropped (B) A schematic depicting the blocked design of the task variants (Simulation, Perception, Control, and Native), as well as the internal composition of a block. (C) A schematic depicting the trial outlines for each of the three variants of interest. The Native variant was not included in the subsequent fMRI analyses and is hence not shown here.

Boards were generated by randomly picking an *x* position, *y* position, length, and angle of rotation value for each of ten planks. We then used the Newton Dynamic library (http://newtondynamics.com), to simulate what would occur if the ball were dropped from its position at the top of a given board. A board was stored if the simulation resulted in the ball falling into one of the two catchers – otherwise, it was discarded, and the process was started over.

In the present study, we sought to determine whether there was neural evidence for the idea that a physical simulation of an object’s trajectory (in this case, the ball) would recruit neural circuits involved in the visual perception of that same trajectory. We hypothesized that if this indeed were the case, then we would observe voxel-wise activity pattern similarity in motion-sensitive regions of the brain between conditions in which participants simulated the ball’s motion trajectory, and conditions in which they perceived the ball’s motion trajectory. To test this hypothesis, we devised three task variants that participants performed in the MRI scanner:

1. *Simulation variant:* On this variant, participants were shown a board and asked to indicate which catcher they thought the ball would land in. This variant thus served as the experimental condition of interest.
2. *Perception variant:* On this variant, the ball dropped on its own as soon as the board appeared, and participants indicated which catcher the ball landed in, after the fact. Participants were instructed to pursue the ball as it fell. This variant thus served as our positive control.
3. *Control variant:* On this variant, participants were shown a board comprised of the same items as the first two variants, except that the planks were more likely to adhere to either a vertical or horizontal orientation. Participants were asked to use this orientation property to inform their response – if the majority of the planks were vertical, they had to press one button, whereas if the majority of the planks were horizontal, they had to press the other. Although the ball was still present on screen during this variant, it was irrelevant to the task and was never dropped into either catcher. Note that the planks in the Simulation and Perception variants could also be horizontal or vertical – they were simply more likely to be so in the Control variant. Thus, for this variant, the visual stimuli were comprised of the same physical objects as in the previous two variants, but the task demands were changed entirely such that no physical interactions needed to be simulated. This allowed us to account for changes in BOLD signal attributable to the visual display (independent of the cognitive process acting on it) as well as eye movements (since all variants permitted free viewing of the scene). This variant served as the negative control.

A schematic depicting the progression of a trial for each of the three variants is shown in Figure 1C. We also had participants perform a fourth variant called the Native variant. This variant was essentially the Simulation and Perception variants combined – participants initially made a judgement about the ball’s trajectory, and once they had responded, we actually dropped the ball for them and let them view whether or not their answer was correct. The function of this variant was largely behavioral – it was necessary to have a condition where participants received immediate visual feedback about their choice so as to ensure that their internal model of the world and its physical properties remained accurate. While participants did perform this variant inside the scanner, the data from this variant was not analyzed for this study (nor were the trial timings optimized for fMRI data analysis). All the data presented comes from the Simulation, Perception, and Control variants.

Participants were pre-trained on all variants, and only progressed to the MRI scanner once they reported feeling comfortable with the demands of the task. Subjects generally reported feeling comfortable with the task after having attempted 10-20 practice trials per variant. Once in the scanner, variants were presented in unified blocks, with approximately 2 minutes of break time provided in between. Each scanning run was made up of one variant block. Each variant block was repeated three times over the course of a session, and variant block order was pseudorandomized. Each block started with a 16 second fixation period during which participants were asked to foveate a fixation spot presented at the center of the screen. Following this fixation period, the block progressed on to 24 task trials. Task variant identity was cued by the color of a fixation spot that was presented at the start of each trial. Trial durations depended on participants’ reaction times and were thus self-paced, although we did impose an eventual 6 second timeout for unusually long trials. Trials had a variable intertrial interval (ITI) of 1-6 seconds, with an average ITI of 2 seconds. Finally, each block ended with another 16 second fixation period during which participants were asked to foveate a fixation spot presented at the center of the screen. A schematic depicting the progression of blocks through a session is shown in Figure 1B. Since each block corresponded to one run and participants performed a total of 3 blocks per task variant, we collected 12 total runs from each participant.

### Classification of Boards

To explicitly relate the properties of the stimuli to physics simulation, we leveraged our control over the embedded Newton Dynamics library to constrain certain properties of the boards. Specifically, we classified boards in the Simulation condition along two relevant dimensions. The first is what we have termed “simulation length”. This dimension refers to the length of the hypothetical simulation required to mentally recreate the ball’s full trajectory for a given board. We use the number of planks hit by the ball as the measure of simulation length because 1) as the number of planks involved increases, the length of the ball’s trajectory also inevitably increases, and 2) each additional plank represents a physical interaction that must be incorporated into the simulation, thereby lengthening it. We divided our boards into two simulation length categories – short (the ball hit 2 planks), or long (the ball hit 4 planks). Examples of short and long simulation length boards are shown in Figures 2A and 2B.

**Figure 2:**
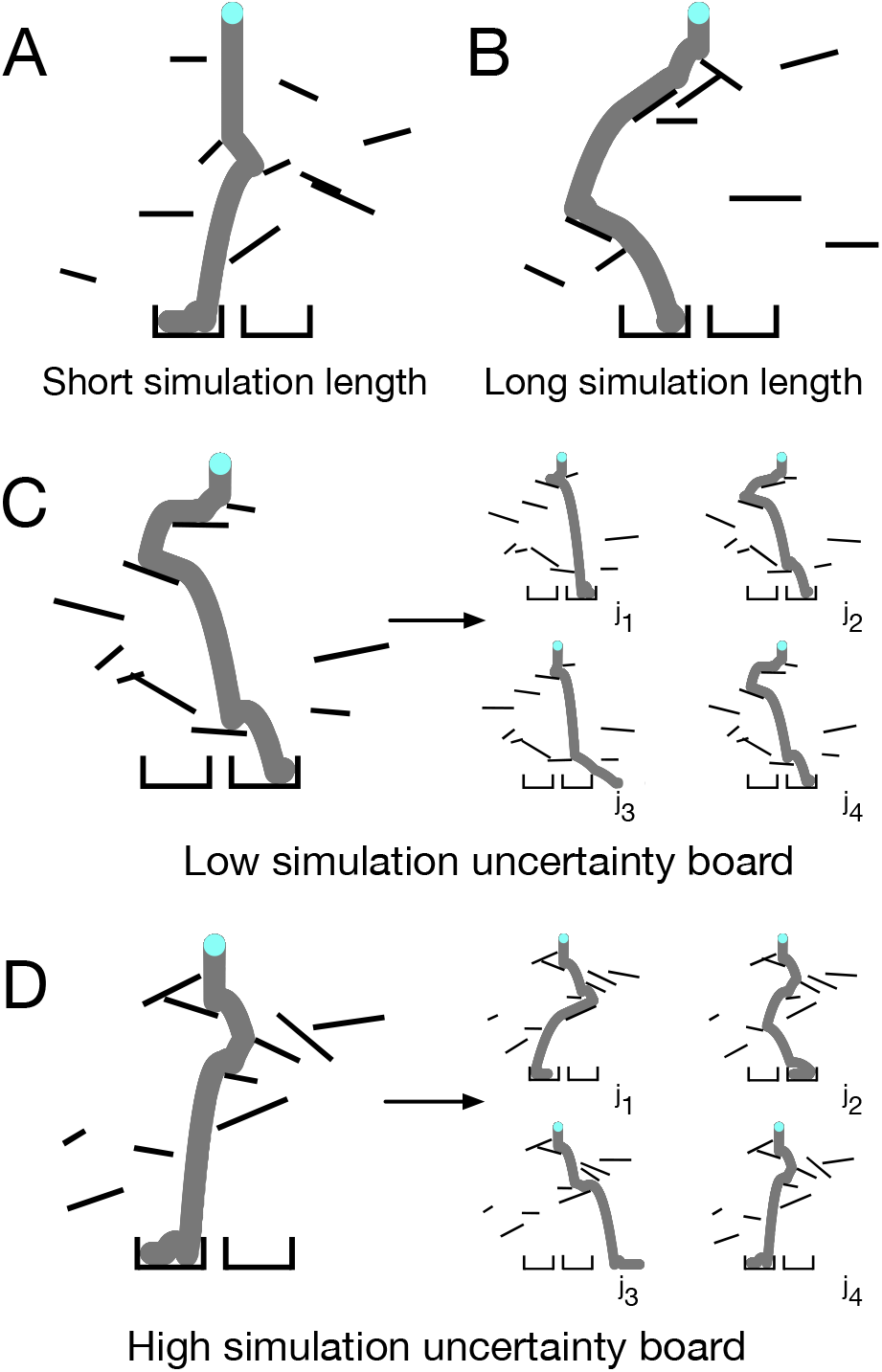
Board Designations for Behavioral Analyses. (A) An example of a board on which the ball only hit two planks. Such boards were classified as having a short simulation length. (B) An example of a board on which the ball hit four planks. Such boards were classified as having a long simulation length. (C) An example of a board where slightly jittering the position of each plank (four jittered examples are shown to the right) had a minimal impact on the ball’s final position. Such boards were classified as having a low simulation uncertainty. (D) An example of a board where slightly jittering the position of each plank (four jittered examples are shown to the right) greatly impacted the ball’s final position. Such boards were classified as having a high simulation uncertainty.

The second dimension that we looked at is what we have termed “simulation uncertainty”. This term refers to the degree of uncertainty involved in simulating a given trajectory. We assigned simulation uncertainty by determining the number of realistic alternate outcomes one might consider within a simulation strategy. Simulation uncertainty was quantified by repeatedly introducing some positional jitter to each plank on a given board, and re-simulating (via our physics engine) the ball’s trajectory on the jittered replicate. This process was carried out offline, 150 times for each board. For some boards, slight jitter of the planks caused major deviations to the ball’s calculated path (relative to the path in the original plank configuration). This meant that a noisy simulation could lead to any one of many plausible outcomes. If the majority of jittered configurations for a given board led to a change in the ball’s final position, that board would be classified as a high simulation uncertainty board. An example of a high simulation uncertainty board is shown in Figure 2D. For other boards, jitter had only a minor effect on the trajectory—the ball generally ended up in the same place. If the majority of jittered configurations for a given board resulted in no change in the ball’s final position, that board would be classified as a low simulation uncertainty board. An example of a low simulation uncertainty board is shown in Figure 2C. A dynamic demonstration of these uncertainty categories is shown in a <u>supplementary video from our previous paper</u> (Ahuja & Sheinberg, 2019). Simulation length and simulation uncertainty categories were counterbalanced, leading to six trials per length and uncertainty combination per block (2 length categories X 2 uncertainty categories X 6 = 24 trials). The Simulation and Perception variant blocks did not use exactly the same set of boards, but the boards in each variant were matched for length and uncertainty.

### fMRI Procedure and Preprocessing

A Siemens 3T PRISMA MRI system with a 64-channel head coil was used for whole-brain imaging. First, a high-resolution T1 weighted multiecho MPRAGE anatomical image was collected for visualization (repetition time, 1900 ms; echo time, 3.02 ms; flip angle, 9°; 160 sagittal slices; 1 × 1 × 1 mm). Functional images were acquired using a fat-saturated gradient-echo echo-planar sequence (TR, 2000 ms; TE, 28 ms; flip angle, 90°; 38 interleaved axial slices; 3 × 3 × 3 mm). Head motion was restricted using padding that surrounded the head. Visual stimuli were displayed on a 24-inch MRI safe screen (Cambridge Research Systems) and viewed through a mirror attached to the head coil. Participants responded using an MR compatible two-button response pad (VPixx Technologies). Preprocessing and analysis of fMRI data were performed using SPM12 (www.fil.ion.ucl.ac.uk/spm). The images were first corrected for differences in slice acquisition timing by resampling slices in time to match the first slice. Next, images were corrected for motion by realigning them to the start of the session using a rigid transformation. Realigned images were then normalized to Montreal Neurological Institute (MNI) stereotaxic space. We opted to not smooth our images and instead preserve as much voxel-unique information as possible because our subsequent analyses focused on individual voxel-level comparisons between conditions.

### Behavioral Analyses

In our earlier experiments, we showed that participants’ reaction times increased commensurately with an increase in simulation length, suggesting that they were engaging in a simulation of the ball’s trajectory. Participants’ accuracy on the task, however, was unaffected by simulation length. Further, we showed that participants’ reaction times were greater and accuracy was lower on high simulation uncertainty boards relative to low simulation uncertainty boards. (Ahuja & Sheinberg, 2019). Here, we sought to replicate these effects from our past experiments to verify that participants’ behavior on the task matched what we have previously presented as evidence in favor of simulation. To that end, we assessed participants’ reaction times and accuracy on the Simulation variant of the task as a function of simulation length and uncertainty.

### General Linear Modeling

To account for neural activity of interest, we modeled our hypothesized BOLD response during the pre-trial period of each variant using a boxcar regressor that spanned the period from stimulus onset till participant response. Because we used a variable duration boxcar model that reflected participants’ reaction times, we were able to appropriately account for the shape of the observed BOLD signal on a trial-by-trial basis, even though each trial lasted for a slightly different amount of time (Grinband et al., 2006; Yarkoni et al., 2009). We modeled the first two trials of each run as nuisance regressors to account for potential noise associated with block initiation and changes in variant identity. Additional nuisance regressors included trials with outlier reaction times, six motion estimates (translation and rotation), and run identity. These were combined with the HRF-convolved task regressors in a design matrix for the entire session. Finally, we used general linear modeling to fit our regressors of interest to the observed BOLD signal and derived beta and t-statistic values by estimating a linear contrast for our task variants relative to an implicit baseline. This analysis was carried out for each of the twelve participants.

### Representational Similarity and Searchlight Analysis

Having derived activity estimates for all variants across all participants, we carried out Representational Similarity Analysis (RSA). In RSA, an activity pattern across a set of voxels for a given condition is treated as the “representation” of that condition in the brain (Kriegeskorte et al., 2008). Representations for various conditions can then be compared to one another to calculate the degree of similarity between them using a metric such as the Euclidean Distance or Spearman correlation (Nili et al., 2014). In the present study, we used the voxel-wise t-statistics derived for each condition (contrasted against baseline) as the activity estimates because they are univariately noise-normalized and have been shown to be more reliable than beta values for RSA (Walther et al., 2015). For the similarity metric, we opted to use a Spearman correlation. Example t-maps from one participant in the Simulation > Baseline, Perception > Baseline, and Control > Baseline contrasts are shown in a motion-sensitive ROI in Figure 3C. We calculated the degree of similarity between the Simulation and Perception conditions (S-P), as well as the Simulation and Control conditions (S-C) for each participant. We then directly compared the observed S-P and S-C similarities to one another (Figure 3D). To reiterate, we hypothesized that if the process of simulation was capable of evoking sensory representations, then the S-P similarity would be greater than the S-C similarity in motion-sensitive areas of the brain.

**Figure 3:**
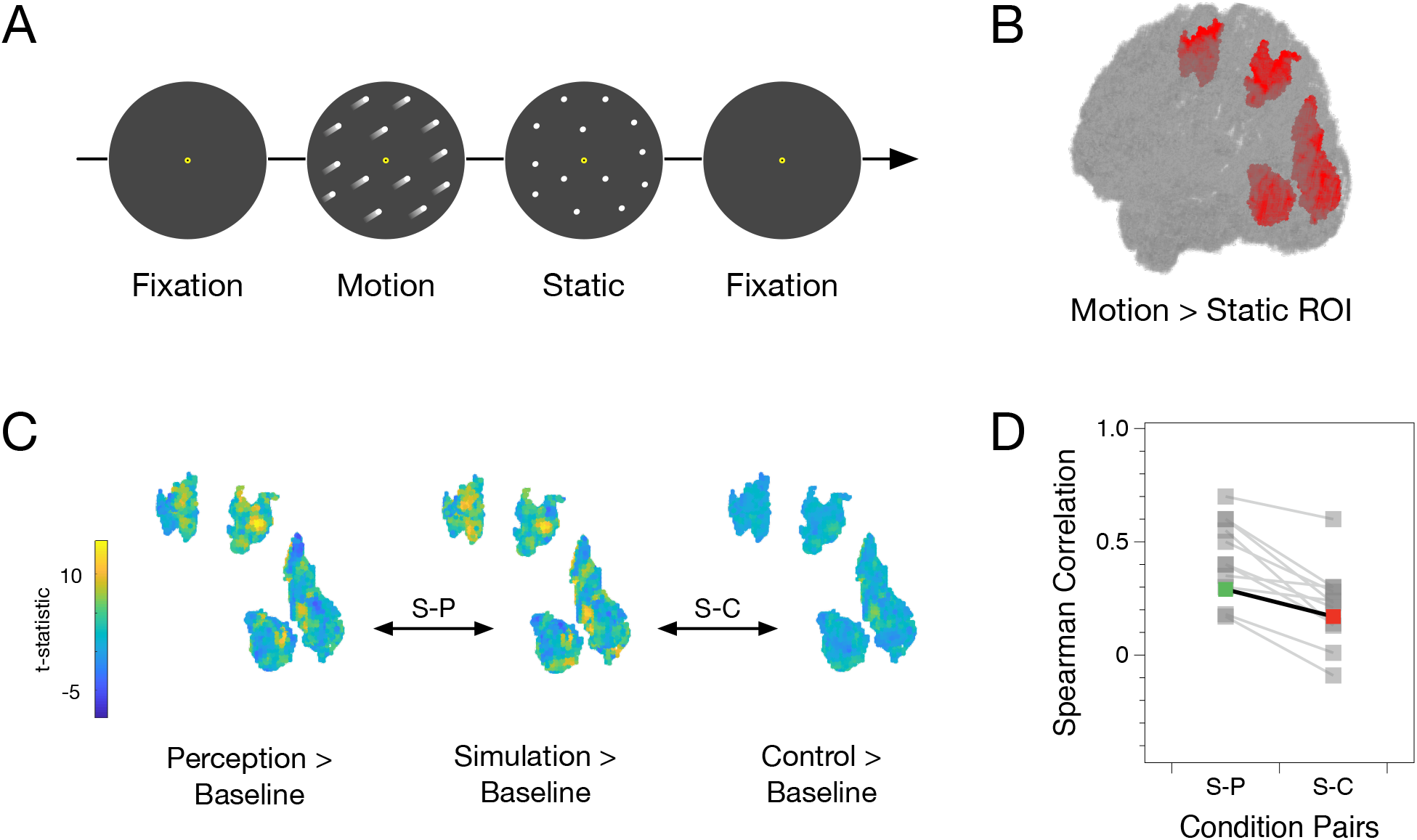
fMRI Analysis Pipeline. (A) A schematic of the motion localizer task. The display alternated between blocks of moving and static dots, flanked by periods of fixation. (B) A hypothetical activation map of motionsensitive voxels derived from a Motion > Static localizer contrast, plotted as a 3D point cloud to demonstrate ROI selection. (C) An example participant’s t-values in the ROI from (B) for each of the three task conditions, contrasted to baseline. Based on these t-maps, we assessed whether the voxel-wise representation in the Simulation condition was more similar to the Perception condition (S-P) or the Control condition (S-C). (D) A line graph showing an example comparison of S-P and S-C similarities. The participant shown in (C) is highlighted in color in (D), and the grey lines represent a hypothetical group effect if the analysis were repeated for all 12 participants.

We also repeated the aforementioned analyses using a searchlight approach. We did this to ensure that any effects that might be observed in the motion-sensitive ROI were in fact specific to those voxels. In a typical searchlight analysis, a cube of voxels is tiled across every possible location in the brain to form several mini-ROIs which can then be analyzed. In the present study, we tiled a 3×3×3 voxel cube over the entire brain for each subject and calculated the representational similarity between conditions at each location. Next, we assessed which loci exhibited a greater S-P similarity compared to S-C similarity in all twelve of our subjects. This approach allowed us to stringently safeguard against the inflated risk of false positives as well as the potential intersubject variability that comes with doing a searchlight analysis (Etzel, Zacks, & Braver, 2013). Finally, we analyzed the brain regions revealed by our searchlight analysis to evaluate whether our effect of interest was indeed specific to these motion-sensitive regions of interest.

## Results

### Behavioral Results

Figure 4A shows participants’ performance on the task across the three variants. Participants were generally very good at all three task variants and performed far better than chance, which was defined as 50% correct (Simulation: 89.8%, [t_11_ = 39.745, p < 0.001]; Perception: 99.6% [t_11_ = 199.23, p < 0.001]; Control: 99.4% [t_11_ = 155.45, p < 0.001]). Subjects’ reaction times in the three variants were significantly different from one another as assessed by a repeated measures analysis of variance (rmANOVA) followed by post-hoc pairwise t-tests with Holm-Bonferroni correction for multiple comparisons (F_1,11_ = 81.85, p < 0.001; Simulation v Control: p < 0.001; Simulation v Perception: p < 0.05; Perception v Control: p < 0.001).

**Figure 4:**
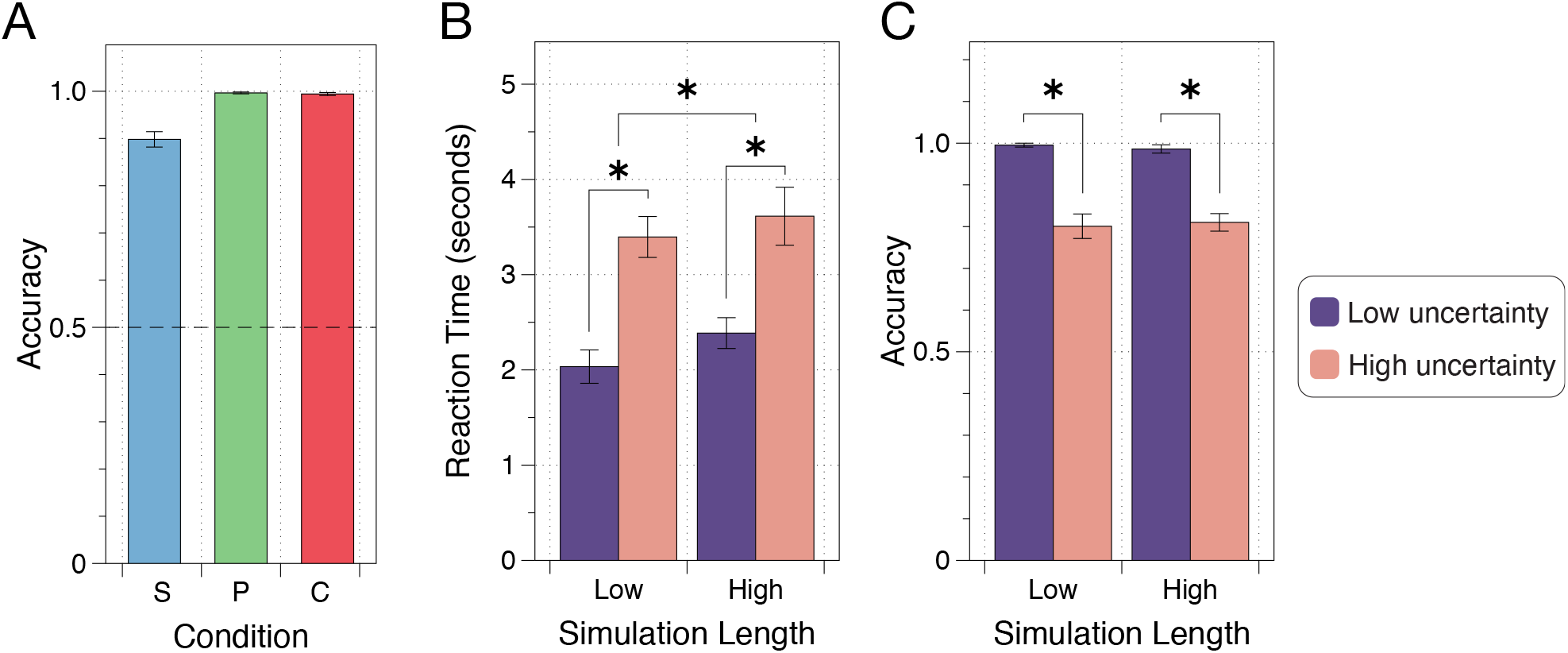
Behavioral Results. (A) Participants’ mean task accuracy across the three variants of interest. Participants were extremely good at all three variants. (B) Participants’ mean reaction times as a function of simulation length and uncertainty. We found that simulation length and simulation uncertainty affected participants’ reaction times on the task. (C) Participants’ average accuracy on the task as a function of simulation length and uncertainty. We found that simulation uncertainty affected participants’ reaction times, whereas simulation length did not. Both of these behavioral effects were previously reported in Ahuja & Sheinberg, 2019. Error bars in all figures represent the standard error of the mean performance for the twelve subjects.

Previously, we showed that people perform the ball fall task via simulation by relating their reaction times and accuracy to two simulation-based metrics – length and uncertainty (Ahuja & Sheinberg, 2019). Specifically, we showed that as simulation length increased, so did participants’ reaction times. We also showed that as simulation uncertainty increased, participants’ reaction times increased, and their accuracy on the task decreased. To ensure that the participants in this study were also engaging in simulation, we repeated this same behavioral analysis on trials from the Simulation variant. Figure 4B shows participants’ average reaction times as a function of simulation length and uncertainty. A twoway repeated measures analysis of variance (rmANOVA) revealed a main effect of both simulation length (F_1,11_ = 6.2, p < 0.05) and simulation uncertainty (F_1,11_ = 51.22, p < 0.001). Figure 4C shows participants’ average accuracy on the task as a function of simulation length and uncertainty. A two-way rmANOVA revealed a main effect of simulation uncertainty (F_1,11_ = 93.78, p < 0.001) but not length (F_1,11_ = 0.04, p = 0.84). We were thus able to recreate the observed effects from our previous work and directly relate participants’ behavior to simulation-based metrics, thereby verifying that participants were indeed engaging in a simulation of the ball’s trajectory during the Simulation variant.

### Localizer Activity

To delineate motion-sensitive voxels that could serve as an ROI for the subsequent analyses, we compared differential neural responses to passive viewing of moving dots versus stationary dots during the localizer task. Several brain regions, such as area MT and the Posterior Parietal Cortex (PPC) were found to be more active during the Motion condition relative to the Static condition in a second-level Motion > Static contrast (Figure 5). Both area MT and PPC have repeatedly been highlighted in past research for their responsivity to moving stimuli, and the same effect is replicated here (Bremmer et al., 2001). Other early visual and premotor areas that have previously been reported to be activated by motion localizer tasks were also observed (Sunaert et al., 1999). Overall, our analysis was successful at identifying motion-sensitive voxels in brain regions that are well-known to be specialized for motion perception. To ensure that the ROI for our subsequent analyses would involve minimal experimenter bias, we used all voxels from the Motion > Static group level contrast (family-wise error [FWE] cluster corrected for multiple comparisons at p < 0.05, extent threshold 187) as our functional ROI. The complete set of activation coordinates for this contrast can be found in Table 1.

**Figure 5:**
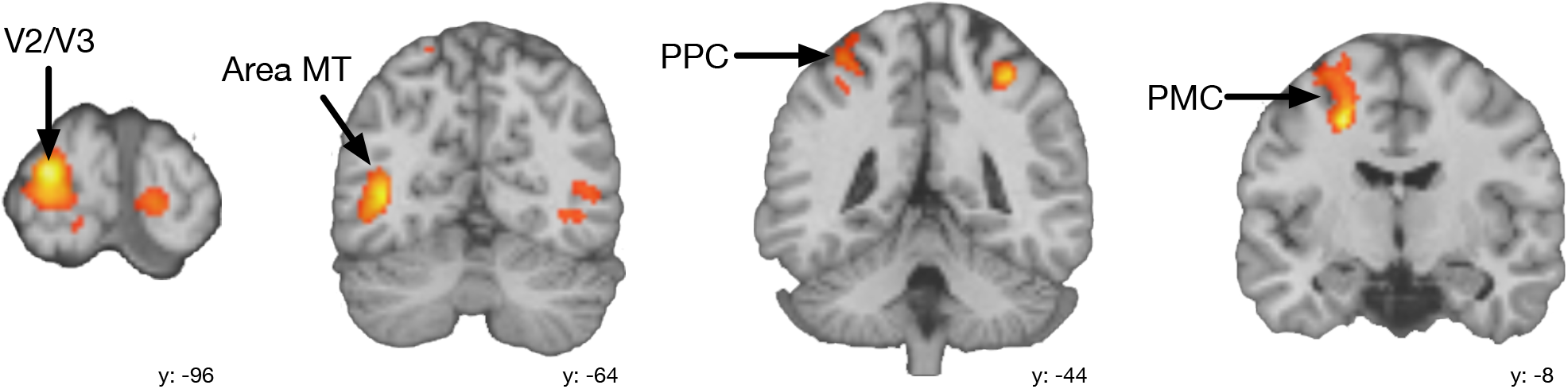
Localizer Results. Activation maps for a second level Motion > Static contrast at a p <0.05 threshold (family-wise error [FWE] cluster corrected for multiple comparisons, extent threshold 187). We observed several canonically motion sensitive regions such as area MT and PPC in this contrast. These voxels were used to define ROIS for subsequent RSA analyses.

**Table 1:** MNI activation coordinates for the Motion > Static contrast from the motion localizer task

| Functional Designation | Anatomical Designation | BA | Peak x | Peak y | Peak z | Peak t-value |
| --- | --- | --- | --- | --- | --- | --- |
| Right V1 | Calcarine fissure | 17 | 16 | -92 | 2 | 4.95 |
| Left V2/V3 | Left middle occipital gyrus | 18 | -22 | -96 | 10 | 8.16 |
| Right V2/V3 | Right middle occipital gyrus | 18 | 26 | -84 | 16 | 5.65 |
| Left V5/MT+ | Left middle temporal gyrus | 19 | -40 | -66 | 6 | 8.1 |
| Right V5/MT+ | Right middle temporal gyrus | 19 | 40 | -70 | 10 | 5.7 |
| Left superior PPC | Left superior parietal gyrus | 7 | -32 | -46 | 60 | 5.79 |
| Right superior PPC | Right inferior parietal gyrus | 7 | 30 | -48 | 50 | 4.32 |
| Left premotor cortex | Left superior frontal gyrus | 6 | -22 | -6 | 60 | 5.03 |

### Task Activity and RSA

Having defined an unbiased functional ROI for motion-responsive voxels in the brain, we turned to a representational similarity analysis of activity estimates in these voxels from the pre-response period of the three task variants. Specifically, we compared the degree of representational similarity between the Simulation and Perception conditions (S-P) as well as the Simulation and Control conditions (S-C). We then compared these S-P and S-C similarity estimates to one another for each participant. There were two potential outcomes of interest, each with a different interpretation. The first possibility was that the Simulation and Control conditions would be more similar to one another in their representations than the Simulation and Perception conditions. Given the sensorimotor properties of each variant and the motion-sensitive ROI, this would not at all be surprising, and might even be expected. After all, the Simulation and Control conditions were both comprised of entirely static displays, and participants made self-directed saccades in each, whereas the Perception condition contained a moving ball in it, and participants largely engaged in guided smooth pursuit of its trajectory. A greater S-C similarity relative to S-P would likely suggest that these voxels faithfully represent the sensory experience of each condition and remain unmodulated by any higher-order cognitive processes or task demands.

The other possibility was that the S-P similarity would be greater than the S-C similarity. If this were the case, it would suggest that despite stimulus-level differences, the cognitive processing engaged in the Simulation condition, i.e., a simulation of the ball’s trajectory, can modulate the activity of motion-sensitive voxels to resemble how these voxels behave when perceiving the ball’s trajectory. In other words, even though the ball’s motion and physical interactions are not being literally perceived, the process of simulating them gives rise to a corresponding sensory representation that is akin to a weak form of perception. In the present study, we predicted that this latter possibility would be the case.

A comparison of the observed S-P and S-C similarity estimates is shown in Figure 6A. The S-P representational similarity was greater than the S-C representational similarity for each of our twelve participants. A paired t-test revealed a significant difference between the S-P and S-C similarities (t_11_ = 5.75, p < 0.001). This finding reflects the second of the two possibilities mentioned earlier and provides evidence for our hypothesis that an internal simulation of physical interactions can give rise to sensory activity in visual areas that look as if one were indeed perceiving these interactions. Next, we wanted to check whether the effect we observed in our multi-region motion-sensitive ROI would also be present in smaller subregions within that ROI. We thus repeated the same RSA in two sub-regions – area MT and PPC – using the bilateral activation clusters outlined in Table 1 as the ROIs. A comparison of the observed S-P and S-C similarity estimates in these ROIs is shown in Figures 6B and 6C. We found that both in Area MT (t_11_ = 7.75, p < 0.001) and PPC (t_11_ = 4.59, p < 0.001), the S-P representational similarity was greater than the S-C representational similarity. This finding shows that the effect shown in Figure 6A is not simply being driven by a small subset of voxels, but that it is present across several motion-responsive functional areas.

**Figure 6:**
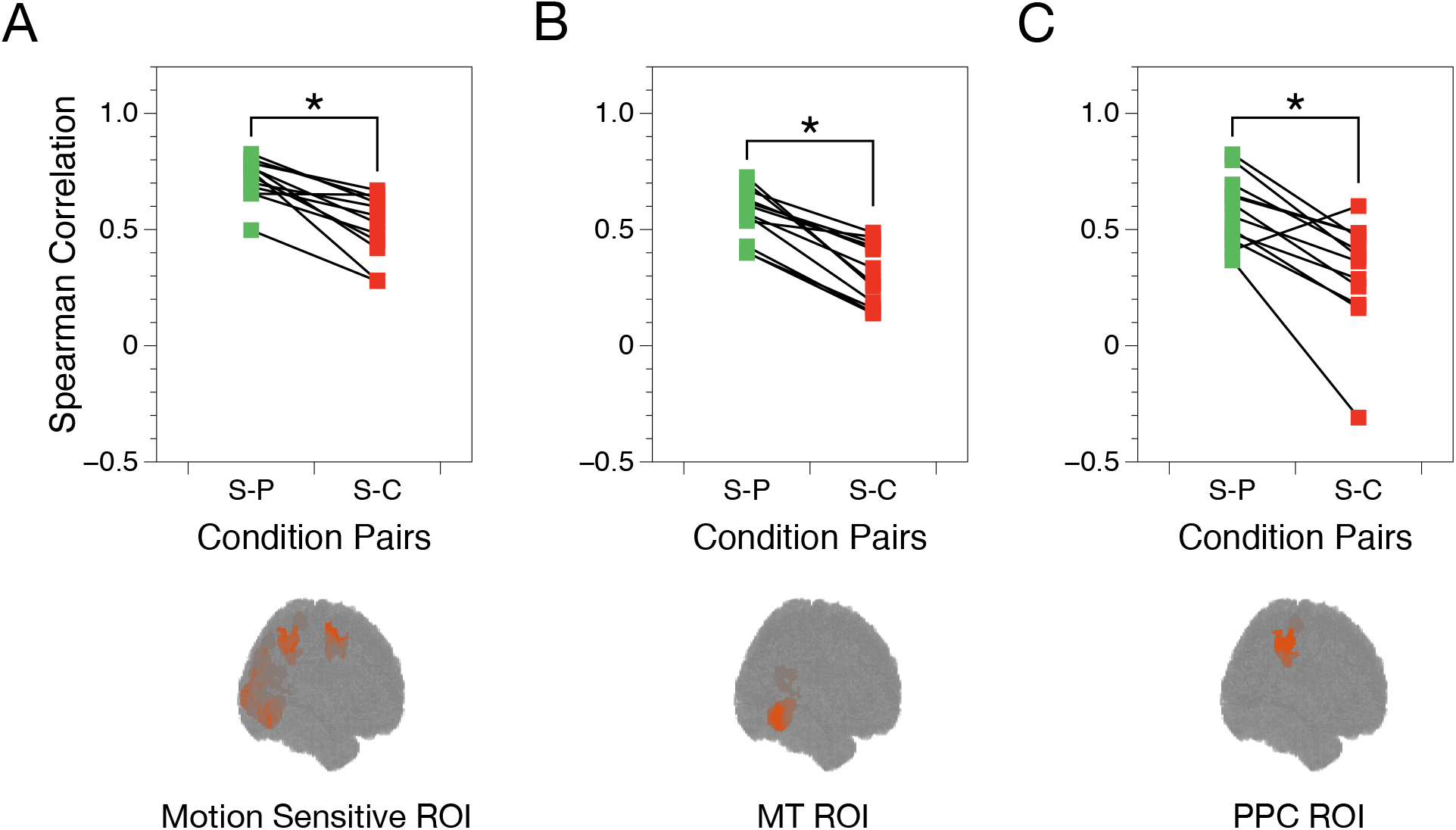
RSA Results. (A) Pairwise comparisons of S-P and S-C representational similarities in a motion-sensitive ROI. Each pair of points represents one participant. We found that for all participants, the representational similarity between the Simulation and Perception conditions was greater (evidenced by a higher Spearman correlation) than the representational similarity between the Simulation and Control conditions (B) The same analysis as in (A), repeated for an MT ROI. (C) The same analysis as in (A), repeated for a PPC ROI.

We further wanted to establish that the observed increase in representational similarity between the Simulation and Perception conditions was not simply a distributed property found all over the entire brain, but that it was specific to our motion-sensitive ROI. We accomplished this by calculating S-P and S-C similarity estimates at every possible 3X3X3 voxel locus in the brain using searchlight analysis (for more details, see Methods). We found that the voxel loci that consistently showed a greater S-P than S-C representational similarity fell largely within the same brain regions that we had already independently isolated using our motion localizer task. These voxel clusters are shown in Figure 7. Table 2 shows a comparison of regions highlighted by the searchlight analysis and the motion localizer. Rows shaded in green represent regions that were identified by both, rows shaded in yellow represent regions that were unilateral in one but bilateral in the other, and rows shaded in blue represent regions that were highlighted by the searchlight only. This result demonstrates that the effect we observed in the motion-sensitive ROIs is indeed quite specific to those voxels.

**Figure 7:**
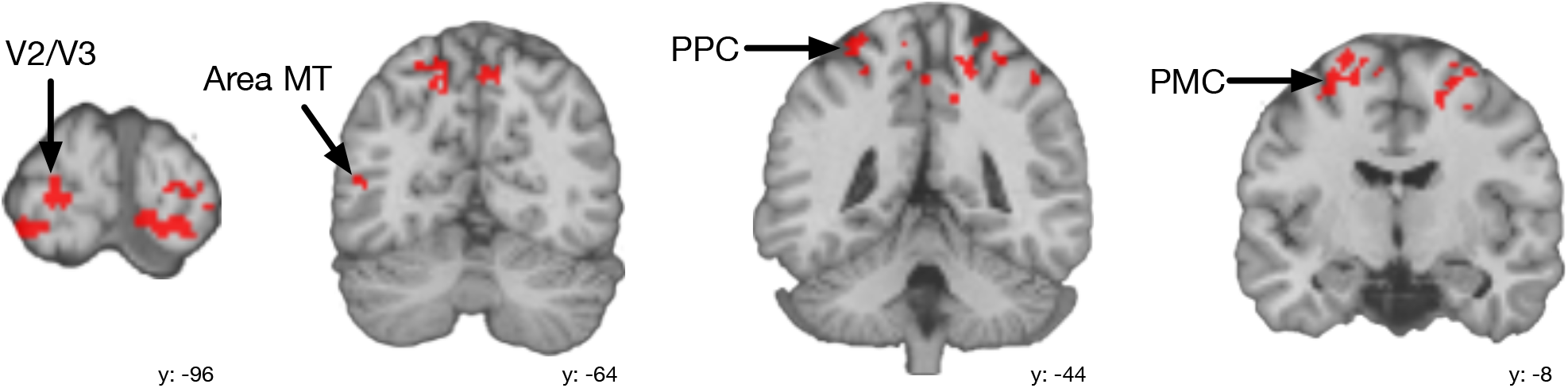
Searchlight Results. Clusters of voxels that were highlighted by a searchlight analysis for consistently exhibiting the main effect from Figure 5. The searchlight largely revealed the same regions as we had previously isolated using the motion localizer task (slices here are the same as in Figure 4).

**Table 2:** Common ROIs highlighted by the motion localizer and the searchlight analysis

| Localizer ROIs | BA | Searchlight ROIs |
| --- | --- | --- |
| Right V1 | 17 | Right V1 |
| Left V2/V3 | 18 | Left V2/V3 |
| Right V2/V3 | 18 | Right V2/V3 |
| Left V5/MT+ | 19 | Left V5/MT+ |
| Right V5/MT+ | 19 |  |
| Left superior PPC | 7 | Left superior PPC |
| Right superior PPC | 7 | Right superior PPC |
| Left premotor cortex | 6 | Left premotor cortex |
|  |  | Right premotor cortex |
|  | 40 | Left inferior PPC |
|  | 39/40 | Right inferior PPC |
|  | 1 | Right primary somatosensory cortex |

## Discussion

Recent studies suggest that simulation is a key cognitive faculty employed to make physics predictions (Ahuja & Sheinberg, 2019; Fischer et al., 2016; Rajalingham et al., 2021). While behavioral and computational evidence supporting this idea is compelling, little is known about the neural mechanisms that underlie such simulations. We theorized that a simulation of a series of events could evoke activity in the brain, akin to how the brain might respond were it to visually perceive the same events. As such, we liken simulation to mental imagery, except with a dynamic internal representation of the external world as opposed to a static one.

In this study, we asked human participants to perform a task in which they had to ascertain the path of a falling ball while in an MRI scanner. We have previously shown (and behaviorally replicated in this study) that participants solve this task via simulation. We found that when participants engaged in a simulation of the ball’s trajectory, motion-sensitive regions of the brain were active, even though no motion was being perceived. Further, this activity bore a high degree of representational similarity to conditions in which participants actually witnessed the ball fall. This finding thus directly complements previous research on mental imagery and rotation, and extends the idea of self-generated sensory representations to dynamic physics simulations (Kaas et al., 2010; Shelton & Pippitt, 2006).

It is worth noting that the opposite perspective has been argued in the past (i.e., that simulations are explicitly non-imagery based) (Hegarty, 2004). This assertion has been based on a few findings. First, past research has shown that when individuals simulate complex mechanical systems (for example, a series of interconnected pulleys), they tend to do so in a piecemeal fashion (Hegarty, 1992). This finding has been used to argue that simulation must not involve holistic visual representations, since if that were the case, the entire scene could be inspected at once and the outcome could be determined without needing to sequentially simulate individual pieces. We would raise the counterpoint that scenes are in fact rarely perceived with uniform salience across one’s visual field, and that complex stimuli are often parsed in a piecemeal fashion via a shifting spotlight of attention (Buschman & Miller, 2010). Given this fact, it is reasonable that a simulated visual representation would also be inspected step-by-step depending on the progression of the corresponding physics simulation. Nonetheless, interesting questions persist about the role of attention in simulation that would be fruitful to explore in future studies. In the present study, the trajectory of the ball and the various plank interactions were inherently sequential in nature, which circumvented this issue entirely.

The second argument against the involvement of visual areas in simulation has to do with reported discrepancies between task performance when participants have their eyes open versus when they have them closed. For instance, it has been shown that when participants are asked to close their eyes and tilt a glass until an imagined amount of water has poured out, they tend to misestimate the exact angle of their own tilt and must usually adjust the tilt angle upon opening their eyes (Schwartz & Black, 1999). This has been interpreted to mean that participants must not have an accurate visual representation of the glass while performing the task with their eyes closed – if they did, they would not need to make adjustments when reopening their eyes. However, there are a few other factors involved in this scenario that must be considered. First, since this specific experiment requires participants to directly interact with the object they are simulating, it is entirely possible that they prioritize motor and proprioceptive information over a visual representation. Just as simulation is not the only strategy one may employ to make physics predictions, it may indeed be that self-generated visual representations only accompany some types of simulations but not others, depending on the task demands and context. Further, it is important to keep in mind that simulated visual representations are unlikely to be perfectly isomorphic and are instead better thought of as useful but crude approximations. As such, it is entirely possible that even if individuals had visually represented the glass in this study, that they would be inclined to make minor refinements when allowed to open their eyes.

Our goal in the present study was to simply assess whether visual areas have any involvement at all in the simulation process. As such, our design doesn’t lend itself to any strong causal conclusions about the contributions of individual brain regions. While this is a limitation of the study, we can still look at the network of regions in which we observed relevant activity for important clues. For instance, we showed that our main similarity effect was present both in area MT, which is known mostly for its role in perception of motion, as well as in PPC, which has been implicated not only in motion perception, but also in spatial reasoning and attentional allocation (Born & Bradley, 2005; Wendelken, 2015). We therefore theorize that PPC might contain the neural apparatus for representing a mental model of the physics within the task, whereas area MT may house the depictive elements of reasoning through it. The two areas may then cooperate as part of a larger network to successfully execute a physics simulation. It is important to note, however, that proponents of mental model theory and mental imagery theory have sought to distinguish the two as separate cognitive frameworks (Sima, Schultheis, & Barkowsky, 2013). For instance, Knauff and Johnson-Laird have argued that mental models are primarily propositional (Knauff & Johnson-Laird, 2002). Kosslyn, on the other hand, places less emphasis on the mental model, and has argued that mental visual depictions are in fact the key piece of the puzzle for reasoning about complicated problems (Kosslyn, Thompson, & Ganis, 2006). We do not see the mental model theory and mental imagery theory as necessarily distinct from one another. In the case of something as complex as a physics simulation, it seems entirely possible that the two theories can converge and that the relevant properties of a scene may be represented in a mental model which is then run with the aid of self-generated visual representations. A theoretical framework that unifies mental models with mental imagery is supported by the neural findings we report in this study. Additional research that specifically probes these theories is necessary to definitively assign causal roles to the various brain regions that likely contribute to the execution of physics simulations.

An important aspect of simulation to consider is the role of eye movements. We felt that it was important to let participants freely view the scene as they attempted to ascertain the ball’s trajectory to ensure that their approach would remain as naturalistic and ecologically valid as possible. To ensure that the effect of eye movements was properly accounted for in our comparisons, we also permitted free viewing of the scene in the Control condition. The Simulation and Control conditions were thus similar to one another at a sensorimotor level – both conditions contained entirely static stimuli as well as self-directed saccades. Conversely, the Perception condition, contained a moving stimulus that participants were instructed to smoothly pursue, making it different from the other two conditions. Despite this fact, the Simulation condition bore a closer neural resemblance to the Perception condition. This finding indicates that the encoded representation in motion-sensitive brain areas is not merely about the eye movements taking place, but rather about the cognitive process that they represent (in this case, simulation). This idea is supported by the theory of deictic coding, which states that eye movements serve to orient and ground cognitive phenomena in the real world (Ballard et al., 1997). Finally, the effect that we highlight here was present even when we constrained the ROI to areas such as area MT which has been shown to be unaffected by saccade-induced retinal motion (Russ et al., 2016).

An alternate potential explanation for the finding we report here is that it is largely driven by the homogenous plank orientations that distinguished the Control variant from the Simulation and Perception variants. This possibility is plausible because regions such as area MT have been shown to contain orientation-selective neurons (Albright, 1984). That said, we observed the similarity effect even in a multi-areal motion-sensitive ROI as shown in Figure 6A. If it were the case that orientation-sensitive MT voxels were exclusively representing the similarity between conditions, it is unlikely that the effect would be present in the large, multi-area ROI. Next, when breaking the larger ROI into its component parts, we also examined anatomical designations outside of just area MT, such as area PPC (Figure 6C). Here too, we clearly observed the effect, even though we are not aware of any literature on PPC neurons responding selectively to oriented bars. The searchlight analysis also failed to raise other brain regions that would lend credence to the orientation hypothesis. Instead, we found that the effect is almost exclusively present in areas highlighted by the motion localizer. Taken together, we believe our findings make the orientation hypothesis unlikely, and that our results are driven by activity representing real and imagined motion.

A final topic worth discussing in a discussion about simulation is the role of experience and familiarity. While the idea of internal visual playback of mental models in the brain is certainly exciting, it also seems likely that implementing simulations in this way would be computationally costly. Intricate and vivid simulations might be useful in some contexts (especially ones that are novel and that permit ample decision time), but they may not always be the optimal approach. For instance, following extensive experience with a certain type of problem, one is likely able to form mental shortcuts which in turn allow for quick, approximate judgements. This fact has been shown to be true of chess players – novices tend to engage in “look ahead” strategies to plan out their moves, whereas grandmasters can make rapid but highly effective moves with only a momentary glance at the board (Calderwood et al., 1988; Gobet & Simon, 1996; Holding & Reynolds, 1982). Given this fact, it is possible that participants with extensive experience on the ball fall task may not necessarily simulate the ball’s trajectory. In such a scenario, subjects’ behavior on the task would likely shift such that it would no longer be well-explained by simulation-based metrics. This idea is supported by the fact that convolutional neural networks that are trained to solve the ball fall task make very different errors than human participants do (Ahuja & Sheinberg, 2019). In regard to neural activity, we would hypothesize that motion-sensitive regions would no longer represent aspects of the ball’s motion, and that activity in these areas would likely more closely resemble what we see in the Control condition. However, more research on this question is warranted before any conclusions can be drawn about whether or not visual representations play a role in facilitating non-simulation based prediction methods.

To conclude, we have presented evidence supporting the idea that when simulating the trajectory of a falling ball, motion-sensitive areas of the brain respond as if the ball’s trajectory were being perceived. This effect is specific to a motion-sensitive ROI and emerges despite differences in oculomotor dynamics during simulation and perception of the ball’s trajectory. These findings suggest that physics simulations evoke observable visual representations. Future research that tackles this question using more causal methods (such as transcranial magnetic stimulation) will be key in further elucidating the exact role of vision in physics simulations.

## Acknowledgements

We would like to acknowledge Dr. Michael Worden, Nadira Yusif-Rodriguez, Dr. Diana Burk, Dr. Shaobo Guan, and Dr. Ryan Miller for their many intellectual contributions to this project. We would also like to thank Dr. Ruobing Xia, Dr. Theresa McKim and Dr. Kati Conen for all their various suggestions and insights.

## References

Ahuja, A., & Sheinberg, D. L. (2019). Behavioral and oculomotor evidence for visual simulation of object movement. Journal of Vision, 19(6), 13–13. https://doi.org/10.1167/19.6.13

Albright, T. D. (1984). Direction and orientation selectivity of neurons in visual area MT of the macaque. Journal of Neurophysiology, 52(6), 1106–1130. doi: 10.1152/jn.1984.52.6.1106

Ballard, D. H., Hayhoe, M. M., Pook, P. K., & Rao, R. P. (1997). Deictic codes for the embodiment of cognition. Behavioral and Brain Sciences, 20(4).

Bates, C. J., Yildirim, I., Tenenbaum, J. B., & Battaglia, P. (2019). Modeling human intuitions about liquid flow with particle-based simulation. PLOS Computational Biology, 15(7), e1007210. https://doi.org/10.1371/journal.pcbi.1007210

Battaglia, P. W., Hamrick, J. B., & Tenenbaum, J. B. (2013). Simulation as an engine of physical scene understanding. Proceedings of the National Academy of Sciences, 110(45), 18327–18332. https://doi.org/10.1073/pnas.1306572110

Born, R. T., & Bradley, D. C. (2005). Structure and Function of Visual Area MT. Annual Review of Neuroscience, 28(1). doi: 10.1146/annurev.neuro.26.041002.131052

Bremmer, F., Schlack, A., Shah, N. J., Zafiris, O., Kubischik, M., Hoffmann, K.-P., Zilles, K., & Fink, G. R. (2001). Polymodal Motion Processing in Posterior Parietal and Premotor Cortex. Neuron, 29(1), 287–296. https://doi.org/10.1016/s0896-6273(01)00198-2

Buschman, T. J., & Miller, E. K. (2010). Shifting the Spotlight of Attention: Evidence for Discrete Computations in Cognition. Frontiers in Human Neuroscience, 4, 194. https://doi.org/10.3389/fnhum.2010.00194

Calderwood, R., Klein, G. A., & Crandall, B. W. (1988). Time Pressure, Skill, and Move Quality in Chess. The American Journal of Psychology, 101(4), 481. https://doi.org/10.2307/1423226

Craik, K. (1943) The Nature of Explanation. Cambridge University Press.

Etzel, J. A., Zacks, J. M., & Braver, T. S. (2013). Searchlight analysis: Promise, pitfalls, and potential. NeuroImage, 78, 261–269. doi: 10.1016/j.neuroimage.2013.03.041

Fischer, J., Mikhael, J. G., Tenenbaum, J. B., & Kanwisher, N. (2016). Functional neuroanatomy of intuitive physical inference. Proceedings of the National Academy of Sciences of the United States of America, 113(34), E5072–81. https://doi.org/10.1073/pnas.1610344113

Gobet, F., & Simon, H. A. (1996). The Roles of Recognition Processes and Look-Ahead Search in Time-Constrained Expert Problem Solving: Evidence From Grand-Master-Level Chess. Psychological Science, 7(1). https://doi.org/10.1111/j.1467-9280.1996.tb00666.x

Grinband, J., Hirsch, J., & Ferrera, V. P. (2006). A Neural Representation of Categorization Uncertainty in the Human Brain. Neuron, 49(5), 757–763. https://doi.org/10.1016/j.neuron.2006.01.032

Hegarty, M. (1992). Mental Animation: Inferring Motion From Static Displays of Mechanical Systems. Journal of Experimental Psychology: Learning, Memory, and Cognition, 18(5), 1084–1102. https://doi.org/10.1037/0278-7393.18.5.1084

Hegarty, M. (2004). Mechanical reasoning by mental simulation. Trends in Cognitive Sciences, 8(6), 280–285. https://doi.org/10.1016/j.tics.2004.04.001

Holding, D. H., & Reynolds, R. I. (1982). Recall or evaluation of chess positions as determinants of chess skill. Memory & Cognition, 10(3), 237–242. https://doi.org/10.3758/BF03197635

Hume, D. (1739). A Treatise on Human Nature (1st ed., L.A Selby-Bigge, ed.). Oxford University Press.

Kaas, A., Weigelt, S., Roebroeck, A., Kohler, A., & Muckli, L. (2010). Imagery of a moving object: the role of occipital cortex and human MT/V5+. NeuroImage, 49(1), 794–804. https://doi.org/10.1016/j.neuroimage.2009.07.055

Klein, I., Dubois, J., Mangin, J.-F., Kherif, F., Flandin, G., Poline, J.-B., Denis, M., Kosslyn, S. M., & Bihan, D. (2004). Retinotopic organization of visual mental images as revealed by functional magnetic resonance imaging. Cognitive Brain Research, 22(1), 26–31. https://doi.org/10.1016/j.cogbrainres.2004.07.006

Knauff, M., & Johnson-Laird, P. N. (2002). Visual imagery can impede reasoning. Memory & Cognition, 30(3), 363–371. doi: 10.3758/bf03194937

Kosslyn, S. M., Ganis, G., & Thompson, W. L. (2001). Neural foundations of imagery. Nature Reviews Neuroscience, 2(9), 635–642. https://doi.org/10.1038/35090055

Kosslyn, S. M., Thompson, W. L., & Alpert, N. M. (1997). Neural Systems Shared by Visual Imagery and Visual Perception: A Positron Emission Tomography Study. NeuroImage, 6(4), 320–334. https://doi.org/10.1006/nimg.1997.0295

Kosslyn, S. M., Thompson, W. L., Kim, I. J., & Alpert, N. M. (1995). Topographical representations of mental images in primary visual cortex. Nature, 378(6556). https://doi.org/10.1038/378496a0

Kosslyn, S. M., Thompson, W. L., & Ganis, G. (2006). The case for mental imagery. Oxford University Press. https://doi.org/10.1093/acprof:oso/9780195179088.001.0001

Kourtzi, Z., & Kanwisher, N. (2000). Activation in Human MT/MST by Static Images with Implied Motion. Journal of Cognitive Neuroscience, 12(1), 48–55. doi: 10.1162/08989290051137594

Kriegeskorte, N., Mur, M., & Bandettini, P. (2008). Representational similarity analysis - connecting the branches of systems neuroscience. Frontiers in Systems Neuroscience, 2, 4. https://doi.org/10.3389/neuro.06.004.2008

Lancaster, J.L., Woldorff, M. G., Parsons, L. M., Liotti, M., Freitas, C. S., Rainey, L., Kochunov, P. V., Nickerson, D., Mikiten, S. A., & Fox, P. T. (2000). Automated Talairach Atlas labels for functional brain mapping. Human Brain Mapping, 10(3), 120–131. https://doi.org/10.1002/1097-0193(200007)10:3<120::aid-hbm30>3.0.co;2-8

Lancaster, J.L., Rainey, L. H., Summerlin, J. L., Freitas, C. S., Fox, P. T., Evans, A. C., Toga, A. W., & Mazziotta, J. C. (1997). Automated labeling of the human brain: A preliminary report on the development and evaluation of a forward-transform method. Human Brain Mapping, 5(4), 238–242. https://doi.org/10.1002/(sici)1097-0193(1997)5:4<238::aid-hbm6>3.0.co;2-4

Lorteije, J. A. M., Barraclough, N. E., Jellema, T., Raemaekers, M., Duijnhouwer, J., Xiao, D., … Wezel, R. J. A. van. (2011). Implied Motion Activation in Cortical Area MT Can Be Explained by Visual Low-level Features. Journal of Cognitive Neuroscience, 23(6), 1533–1548. doi: 10.1162/jocn.2010.21533

Maldjian, J. A., Laurienti, P. J., & Burdette, J. H. (2004). Precentral gyrus discrepancy in electronic versions of the Talairach atlas. NeuroImage, 21(1), 450–455. https://doi.org/10.1016/j.neuroimage.2003.09.032

Maldjian, J. A., Laurienti, P. J., Kraft, R. A., & Burdette, J. H. (2003). An automated method for neuroanatomic and cytoarchitectonic atlas-based interrogation of fMRI data sets. NeuroImage, 19(3), 1233–1239. https://doi.org/10.1016/s1053-8119(03)00169-1

Nili, H., Wingfield, C., Walther, A., Su, L., Marslen-Wilson, W., & Kriegeskorte, N. (2014). A Toolbox for Representational Similarity Analysis. PLoS Computational Biology, 10(4), e1003553. https://doi.org/10.1371/journal.pcbi.1003553

Pearson, J. (2019). The human imagination: the cognitive neuroscience of visual mental imagery. Nature Reviews. Neuroscience, 20(10), 624–634. doi: 10.1038/s41583-019-0202-9

Rajalingham, R., Piccato, A., & Jazayeri, M. (2021). The role of mental simulation in primate physical inference abilities. BioRxiv, 2021.01.14.426741. https://doi.org/10.1101/2021.01.14.426741

Russ, B. E., Kaneko, T., Saleem, K. S., Berman, R. A., & Leopold, D. A. (2016). Distinct fMRI Responses to Self-Induced versus Stimulus Motion during Free Viewing in the Macaque. The Journal of Neuroscience, 36(37). https://doi.org/10.1523/jneurosci.1152-16.2016

Schwartz, D. L., & Black, T. (1999). Inferences Through Imagined Actions: Knowing by Simulated Doing. Journal of Experimental Psychology: Learning, Memory, and Cognition, 25(1), 116–136. https://doi.org/10.1037/0278-7393.25.1.116

Shelton, A. L., & Pippitt, H. A. (2006). Motion in the mind’s eye: Comparing mental and visual rotation. Cognitive, Affective, & Behavioral Neuroscience, 6(4), 323–332. https://doi.org/10.3758/cabn.6.4.323

Sima, J. F., Schultheis, H., & Barkowsky, T. (2013). Differences between Spatial and Visual Mental Representations. Frontiers in Psychology, 4, 240. doi: 10.3389/fpsyg.2013.00240

Smith, A. (1759). A Theory of Moral Sentiments. Clarendon.

Sunaert, S., Hecke, P., Marchal, G., & Orban, G. (1999). Motion-responsive regions of the human brain. Experimental Brain Research, 127(4), 355–370. https://doi.org/10.1007/s002210050804

Ullman, T. D., Spelke, E., Battaglia, P., & Tenenbaum, J. B. (2017). Mind Games: Game Engines as an Architecture for Intuitive Physics. Trends in Cognitive Sciences, 21(9), 649–665. https://doi.org/10.1016/j.tics.2017.05.012

Vernet, M., Quentin, R., Chanes, L., Mitsumasu, A., & Valero-Cabré, A. (2014). Frontal eye field, where art thou? Anatomy, function, and non-invasive manipulation of frontal regions involved in eye movements and associated cognitive operations. Frontiers in Integrative Neuroscience, 8, 66. https://doi.org/10.3389/fnint.2014.00066

Walther, A., Nili, H., Ejaz, N., Alink, A., Kriegeskorte, N., & Diedrichsen, J. (2015). Reliability of dissimilarity measures for multi-voxel pattern analysis. NeuroImage, 137, 188–200. https://doi.org/10.1016/j.neuroimage.2015.12.012

Wendelken, C. (2015). Meta-analysis: how does posterior parietal cortex contribute to reasoning? Frontiers in Human Neuroscience, 8, 1042. doi: 10.3389/fnhum.2014.01042

Yarkoni, T., Barch, D. M., Gray, J. R., Conturo, T. E., & Braver, T. S. (2009). BOLD correlates of trial-by-trial reaction time variability in gray and white matter: a multi-study fMRI analysis. PloS One, 4(1), e4257. https://doi.org/10.1371/journal.pone.0004257

